# HIV-1 RNA genomes initiate host protein packaging in the cytosol independently of Gag capsid proteins

**DOI:** 10.1101/846105

**Authors:** Jordan T. Becker, Edward L. Evans, Bayleigh E. Benner, Stephanie L. Pritzl, Laura E. Smith, Andra E. Bates, Nathan M. Sherer

**Affiliations:** McArdle Laboratory for Cancer Research, Institute for Molecular Virology, and Carbone Cancer Center, University of Wisconsin-Madison, Madison, WI, 53706, US

## Abstract

HIV-1 RNA genomes interact with diverse RNA binding proteins (RBPs) in the cytoplasm including antiviral factor APOBEC3G (A3G) that, in the absence of viral Vif proteins, is packaged into virions. Where and when HIV-1-A3G interactions are initiated for packaging inside the cell is unknown, and the relative contributions of genome vs. Gag capsid proteins to this process remains controversial. Here we visualized A3G responses to HIV-1 infection over an entire replication cycle using long-term (up to 72 h) live single cell imaging. We show that Vif-deficient HIV-1 dramatically shifts A3G and a second RNA surveillance factor, MOV10, from the cytoplasm to virus particle assembly sites with little to no net discernible effects on general markers of cytoplasmic processing bodies (DCP1A), stress granules (TIA-1), or a marker of the nonsense-mediated decay machinery (UPF1). Using a new live cell RNA-protein interaction assay based on RNA tethering (the <u>i</u>n-<u>c</u>ell RNA-protein <u>i</u>nteraction <u>p</u>rotocol, or IC-IP), we provide evidence that A3G- and MOV10- genome interactions are selective, strong, occur in presence or absence of Gag, and are initiated in the cytosol soon if not immediately after genome nuclear export. Finally, although Gag is sufficient to package A3G into virions even in the absence of genomes, single virion imaging indicates that selective A3G-genome interactions promote much more consistent per virion delivery of A3G to assembly sites. Collectively, these studies suggest a paradigm for early, strong, and persistent cytosolic detection of select HIV-1 RNA signatures by A3G, MOV10 and other host RBPs that are enriched in virions.

**IMPORTANCE:** Host-pathogen interactions determine the success of viral replication. While extensive work has identified many interactions between HIV-1 and cellular factors, our understanding of where these interactions in cells occur during the course of infection is incomplete. Here, we show that multiple RNA-binding proteins (including the antiviral restriction factor, APOBEC3G, and MOV10 helicase) bind HIV-1 RNA genomes in the cytoplasm and co-traffic with them into progeny virions. Furthermore, we show that these interactions with HIV-1 RNA occur in the absence of Gag and are sufficiently strong to recruit these to otherwise non-native subcellular locales. Together, these data begin to illuminate the intracellular trafficking pathways shared by host RNA binding proteins and the viral RNAs they preferentially bind.

## INTRODUCTION

The human immunodeficiency virus type 1 (HIV-1) hijacks a diverse set of host cellular RNA binding proteins (RBPs) to carry out viral RNA transcription, nuclear export, translation, and trafficking (1–4). Select host RBPs are packaged into virions and exhibit antiviral properties, with the best-characterized example being members of the Apolipoprotein B mRNA editing enzyme, catalytic polypeptide-like 3 (APOBEC3) family. A subset of APOBEC3 proteins (A3F, A3G, and A3H) can be packaged into virions to abolish infectivity by deaminating cytidines on the nascent minus-sense DNA strand of the viral genome, thereby generating G-to-A mutations in the DNA provirus (5–8). During productive infection, however, APOBEC3 proteins are counteracted by viral infectivity factor (Vif) proteins that facilitate their proteasome-mediated degradation prior to the onset of virus particle production (9–11). Additional host factors that are unaffected by Vif are packaged into virions, some of which, *e.g.*, RNA helicases MOV10 and UPF1, have been implicated as regulatory factors influencing virion infectivity (12–16).

How APOBEC proteins evolved to ensure encapsidation into virions remains a poorly understood aspect of the cell-intrinsic host defense. HIV-1 virions are, in essence, transmissible ribonucleoprotein (RNP) complexes, with their assembly driven by viral Gag capsid proteins that multimerize on an RNA scaffold consisting of host-derived RNA molecules and two dimerized viral RNA genomes, with Gag-RNA binding mediated by Gag’s C-terminal Nucleocapsid (NC) domain [reviewed in (17, 18)]. In the absence of viral genomes, Gag expression is sufficient to drive formation of non-infectious virus-like particles containing cellular RNAs, packaged in proportion to their relative abundance in the cell (19, 20). A3G’s incorporation into virions has been shown, for most cell types, to be both RNA- and NC-dependent (21–26)(22, 27). However, the relative contributions of viral genomes vs. host RNAs to this process remains controversial. On the one hand, A3G can incorporated into virus particles through promiscuous interactions with host RNA even when packageable genomes are not expressed (23, 25, 27). On the other hand, A3G incorporation levels can be enhanced by genomes (16, 21); and A3G exhibits selective RNA-binding characteristics (25, 28) including a reported preference for G-rich segments of the HIV-1 genome (26). One or more additional host factors may contribute to A3G packaging including the 7SL non-coding RNA component of the signal recognition particle (29–32).

Where and when A3G and other packaged RBPs interface with Gag and/or genomes in the cell has also been under investigation for some time, with conflicting results. At steady-state, A3G is distributed throughout the cytosol (the aqueous phase of the cytoplasm) and accumulates to high levels at non-membranous cytoplasmic sites of mRNA decay known as processing bodies (PBs) (33, 34). Cytosolic A3G will also rapidly re-localize to sites of translational repression known as stress granules (SGs) in response to heat shock or oxidative stress (33, 34). Although early studies suggested a functional link between PBs and A3G’s antiviral activity (34–36); we have not observed viral genomes and Gag to be overtly associated with P bodies (37–39), and dissolution of these structures by depleting P body structural components has no effect on HIV-1 replication (37, 40). Because HIV-1 Gag and genomes fill the cytoplasm in a non-localized fashion prior to virion assembly (38, 39, 41, 42), we predicted that the virus-host interactions precipitating A3G packaging were most likely to occur ubiquitously throughout the fluid phase of the cytosol. However, such a scenario has yet to be formally demonstrated.

To gain greater understanding of the subcellular dynamics of HIV-1 RNA trafficking, virion production, and host cellular response to infection, herein we carried out a comprehensive analysis of the coordinated behaviors of HIV-1 RNA genomes, Gag, A3G, and markers of PBs, SGs, and the NMD machinery in single cells using single cell video microscopy, in conjunction with a new tethering-based intracellular RNA-protein interaction assay (the <u>i</u>n-<u>c</u>ell RNA-protein <u>i</u>nteraction <u>p</u>rotocol, or IC-IP. Collectively, our results indicate that viral genomes exhibit markedly strong and selective interactions with A3G and at least one additional host RBP, MOV10; that these interactions are Gag-independent; and that these interactions occur throughout the cytosol and are initiated soon if not immediately after viral genome nuclear export.

## RESULTS

### Single cell imaging using RBP biosensor cell lines to track HIV-1 replication and detect host protein packaging

To study real-time RBP responses to HIV-1 in single cells over an entire round of viral replication, we generated biosensor HeLa cell lines stably expressing fluorescent protein-tagged versions of A3G (YFP-A3G) (Fig. 1A) and, as comparators, cell lines constitutively expressing YFP-MOV10 (Fig. 1I), YFP-UPF1 (a component of the NMD machinery) (Fig. 1I), TIA-1-YFP (a marker of SGs) (Fig. 2A) or CFP-DCP1A (a marker of PBs) (Fig. 2E) (33, 63). To address the role(s) of Vif, cells were infected with either Vif-competent (Vif+) HIV-1_NL4-3_ CFP reporter virus or viruses rendered Vif-minus (Vifxx) due to the insertion of two stop codons in the *vif* reading frame (see cartoon depiction in Fig. 1B) (mutation based on ref. (52)). In the absence of infection, YFP-A3G was localized to the cytoplasm and accumulated in bright PBs (Fig. 1A, left panel), with induction of oxidative stress using 250 µM sodium arsenite (Ars) causing re-localization of YFP-A3G to SGs, as anticipated (64)(Fig. 1A, right panel). We also confirmed that our YFP-A3G fusion protein was packaged into Vifxx virus particles and retained antiviral activity using single-round assembly and infectivity assays (Fig. 1C).

**Figure 1.**
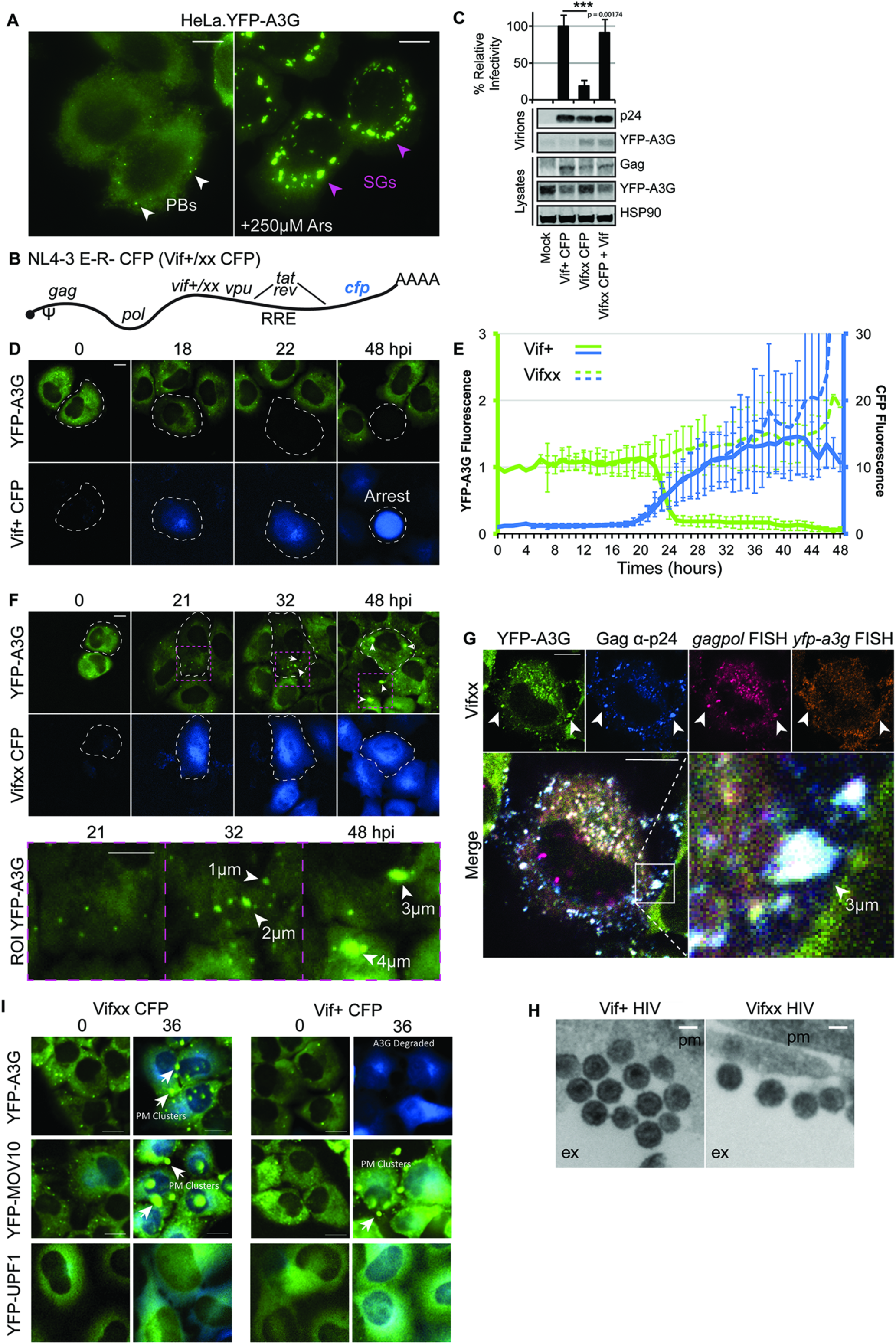
Tracking the HIV-1 replication cycle using a YFP-A3G biosensor cell line. **(A)** HeLa.YFP-A3G cells showing diffuse and punctate (processing bodies, PB, white arrows) distribution of YFP-A3G at steady-state (left) and accumulation into stress granules (SGs, magenta arrows) following 250 µM arsenite treatment (right, Ars). **(B)** Schematic of Vif-competent (Vif+) or Vif-deficient (Vifxx) HIV-1 NL4-3 Env-, Vpr-, CFP reporter virus genome. **(C)** Single-round infection confirms YFP-A3G-mediated restriction of HIV-1 in the absence of Vif. 293T.YFP-A3G cells were transfected to generate the indicated HIV-1 reporter viruses with or without co-expression of Vif-mCherry. Cells and virus particles (*n*=3) were harvested at 48 h post-transfection and analyzed by quantitative immunoblot using anti-p24^Gag^, anti-YFP, and anti-HSP90 (loading control) antisera. **(D)** Representative images from long-term time-lapse imaging showing degradation of YFP-A3G after HIV-1 Vif+ CFP infection and Vif-dependent G2/M arrest at later (∼48h) time points. Figure corresponds to Supplementary Movie 1. **(E)** Quantification of movies (IntDen) represented in (D) (Vif+, solid lines) and (Vifxx, dashed lines). Cells from 3 independent experiments (*n*=33) with ≥16 hours per cell (1 image/hour). **(F)** Representative images from long-term time-lapse imaging of YFP-A3G upon HIV-1 Vifxx CFP infection. Magenta boxes highlight regions of interest (ROIs) shown below and white arrows indicate YFP-A3G clustering at PM. Figure corresponds to Supplementary Movie 2. **(G)** Fixed cell images show Gag and *gagpol* mRNA (viral RNA genomes) accumulating with YFP-A3G at the cell periphery. White arrows highlight cell-peripheral clusters of YFP-A3G, Gag, and genome. (**H)** TSEM images showing virus particles clustering in the extracellular space after budding from HeLa.YFP-A3G cells infected with either Vif+ or Vifxx viruses. All scale bars in fluorescent images = 10µm and in TSEM = 100nm. Error bars represent standard deviation of the mean. **(I)** Representative images from long-term time-lapse imaging showing YFP-A3G, YFP-MOV10, and YFP-UPF1 upon HIV-1 Vifxx and Vif+ infection. Arrows highlight PM clusters. Figure corresponds to Movie S3.

**Figure 2.**
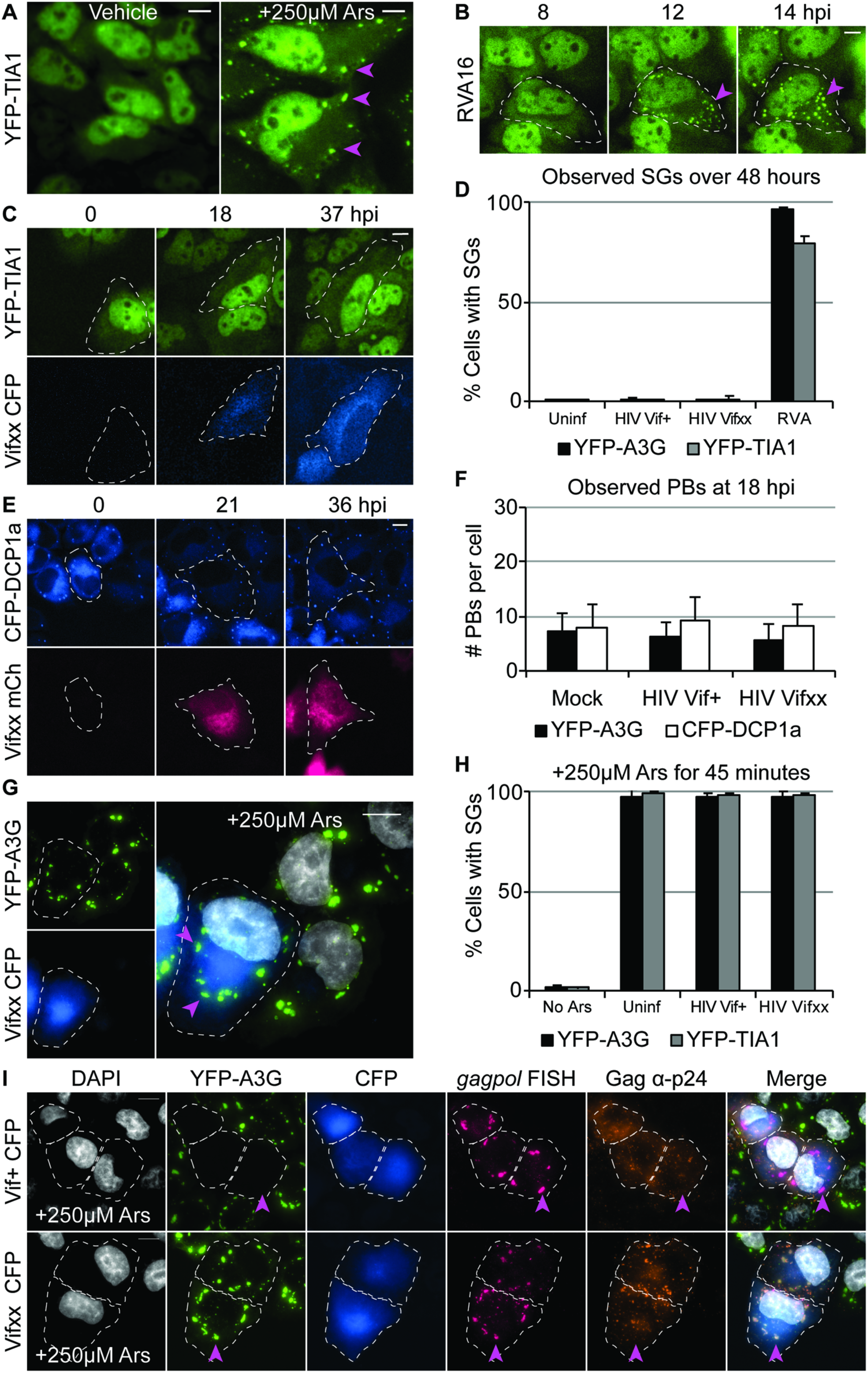
HIV-1 infection has little to no impact on P-bodies or stress granules. **(A)** HeLa.YFP-TIA-1 cells showing steady-state distribution (H_2_0 vehicle, left) and accumulation into SGs (magenta arrows) following arsenite treatment (right, Ars). **(B)** Representative images from time-lapse imaging showing induction of SGs during infection by RVA16 (MOI=10). Figure corresponds to Movie S4. **(C)** Time-lapse images showing no effect on YFP-TIA-1 distribution following Vifxx CFP infection, quantified in **(D)** for >400 HeLa.YFP-A3G (black bars) and HeLa.YFP-TIA-1 (gray bars) cells per condition, combined from three independent experiments. **(E)** Time-lapse images showing no effect on CFP-DCP1A following Vifxx mCh infection, quantified in **(F)** for 30 cells from 3 movies at ∼18 hpi (YFP-A3G, black bars; CFP-DCP1A, white bars). **(G)** HeLa.YFP-A3G infected with Vifxx CFP and treated with Ars to induce SGs at 24 hpi and quantified in **(H)** for >100 infected cells from three independent experiments. Magenta arrows highlight SGs present in infected cells. **(I)** Simultaneous FISH/IF to detect *gagpol* mRNA and Gag show HIV RNA genomes accumulating in SGs in HIV infected cells following Ars treatment. Magenta arrows highlight SGs present in infected cells. All scale bars in fluorescent images = 10µm. Error bars represent standard deviation of the mean.

Hela.YFP-A3G cells were infected with Vif+ virus at a MOI of <1 so that infected and uninfected cells would be visualized side-by-side over an entire round of replication, requiring >48 hours of continuous single cell imaging. Viral gene expression (based on the CFP reporter) was initiated 16-20 hours post-infection (hpi), followed by gradual loss of YFP-A3G from 20-25 hpi consistent with Vif-driven degradation (decay rate of 24.5% per h +/− 6.0%; n=33; Fig. 1D, Movie S1, and quantification in Fig. 1E). YFP-A3G was depleted from both diffuse cytosolic and PB pools simultaneously, with levels suppressed to near background with little to no discernible fluctuations to Vif activity after 24h (Fig. 1D and Fig. 1E showing kinetics over the entire 48h time course). CFP+ cells ultimately became rounded after 40 hpi, consistent with Vif-induced cell cycle arrest (65, 66) (Fig. 1D and Movie S1, see 48 h time point).

We next infected HeLa.YFP-A3G cells with Vif-deleted (Vifxx) reporter virus. As expected, YFP-A3G expression was maintained in the cytoplasm and at PBs throughout the entire course of infection, and cell cycle arrest was not observed (Fig. 1F, quantification in Fig. 1E and Movie S2). Strikingly, however, in infected cells much of the YFP-A3G signal coalesced into large, up to 4 µm diameter (based on current resolution) clusters at 30-48 hpi (Fig. 1F, white arrows highlight growing clusters); unlikely to be PBs or SGs due to their proximity to the plasma membrane and real-time observations of cluster release and transfer to neighboring cells (Fig. 1F, 32h time point, and Movie S2). YFP-A3G clusters stained positive for both HIV-1 Gag/Gag-Pol and viral genome (unspliced RNA encoding the *gagpol* reading frame) but not *yfp-a3g* mRNA (negative control) detected using a 4-color combined anti-p24^Gag^ immunofluorescence / RNA fluorescence in situ hybridization (IF/FISH) protocol, thus indicating that the clusters most likely represented YFP-A3G recruited to sites of virus particle assembly (Fig. 1G). Thin section electron microscopy confirmed the presence of clusters of virus particles at the surface of HeLa.YFP-A3G cells at 48 hpi for both Vif+ and Vifxx conditions, consistent with this hypothesis (Fig. 1H).

For comparison, we also monitored Vif+ and Vifxx infection in cells expressing YFP-MOV10, a superfamily 1 (SF1) RNA helicase and PB protein that is also efficiently packaged into virions and interacts with RNA but, unlike A3G, is not targeted for proteosomal degradation by Vif (13, 14); and cells stably expressing another YFP-tagged SF1 helicase, UPF1, that plays a key role in NMD and is also packaged into virions (12, 67) (Fig. 1I, Fig. S1A, and Movie S3). We observed very similar effects of HIV-1 infection on YFP-MOV10 as for YFP-A3G, with large amounts of signal transferred to large plasma membrane clusters at later time points post-infection; both in the presence or absence of Vif (Figs. 1I, S1B, S1C, and Movie S3). By contrast, YFP-UPF1, that exhibits a diffuse distribution in the cytoplasm, was largely unaffected by HIV-1 (Figs. 1I, S1C, and Movie S3); consistent with it being less efficiently packaged into virions relative to YFP-MOV10 (Fig. S1A).

Taken together, these experiments validated both YFP-A3G and YFP-MOV10 as proxy biosensors for detecting and monitoring HIV-1 infection, with A3G providing for single cell measurements of Vif-induced A3G degradation and cell cycle arrest; and both YFP-A3G (in the absence of Vif) and YFP-MOV10 allowing for direct detection of HIV-1 virus particle assembly and host protein packaging as it occurs.

### HIV-1 replication has no overall effect on processing bodies or stress granules

Because A3G and MOV10 are enriched in SGs and PBs (33, 34, 37, 40), to address specificity and determine if there are transitional effects of HIV-1 on SGs and PBs over time we next studied biosensor cell lines expressing YFP-TIA-1 or CFP-DCP1A (Fig. 2); validated markers of SGs and PBs, respectively [reviewed in (63)]. We first confirmed that the YFP-TIA-1 “stress” biosensor formed SGs in response to sodium arsenite treatment (Ars) (Fig. 2A, right panel) or in response to infection by using rhinovirus A16 (RVA16), a positive-strand RNA virus we had studied previously (68) and discovered to cause rampant SG formation as early as 4h post-infection (Fig. 2B and Movie S4). CFP-DCP1A was enriched in bright punctae throughout the cytoplasm consistent with PBs, as expected (Fig. 2E).

Unlike for RVA16, neither Vif+ nor Vifxx HIV-1 triggered SG formation in either YFP-TIA-1 or YFP-A3G cells over 48h of continuous imaging, a time window sufficient to encompass an entire round of viral replication (Fig. 2C with quantification in Fig. 2D). HIV-1 also had no discernible effects on PB number or morphology in CFP-DCP1A biosensor cells; with PB number being remarkably consistent and stable for individual cells under all conditions (typically 6-9 PBs per cell, Fig. 2E with quantification in Fig. 2F). Unlike YFP-A3G and YFP-MOV10, neither YFP-TIA-1 or CFP-CDP1A were shifted to the plasma membrane over the imaging time course. Based on these observations, we conclude that despite HIV-1’s marked effects on YFP-A3G and YFP-MOV10 as shown in Figure 1, net modulation of PBs or induction of SGs are unlikely to be intrinsic features of the HIV-1 replication cycle.

HIV-1 Gag has been reported to inhibit SG formation (69–72), so that we also tested the effects of treating HeLa.YFP-A3G or HeLa.YFP-TIA-1 biosensor cells with sodium arsenite at 31 hpi when NL4-3 Gag expression in HeLa cells is relatively high (*e.g.*, see ref.(66)). Contrary to our expectations, both YFP-A3G and YFP-TIA-1 accumulated in SGs in Ars-treated cells with or without infection, for both Vif+ and Vifxx viruses (Fig. 2G with quantification in Fig. 2H). To localize viral genomes under these conditions, we again performed combined *gagpol* FISH and Gag IF, finding the bulk of the HIV-1 *gagpol* signal markedly co-localized with YFP-A3G in Ars-induced SGs (Fig. 2I, compare YFP-A3G panels to *gagpol* FISH panels). By contrast, the cytoplasmic pool of Gag was largely excluded from these complexes (Fig. 2I, compare YFP-A3G panels to Gag anti-p24 panels). SGs are induced after kinase R-mediated phosphorylation of translation initiation factor EIF2A, triggering aggregation of scaffolding proteins TIA-1, TIAR, and G3BP1 in complex with mRNAs bound to translation initiation factors and polysome-associated proteins [reviewed in (63, 73)]. Accordingly, marked co-localization of YFP-A3G viral unspliced RNA in SGs would be consistent with A3G co-trafficking with genomes and/or *gagpol* mRNAs in a cytosolic phase that contains free polysomes.

### Evidence for strong genome-A3G interactions in the cytosol based on FRAP

In order to be able to directly track genomes and Gag in conjunction with YFP-A3G and YFP-MOV10, we next engineered two-color visible HIV-1 viruses expressing Gag-CFP from “self-tagging” RNA genomes bearing 24 copies of the MS2 RNA binding loop (located in the *gag-pol* open reading frame) and encoding an RFP-tagged MS2 bacteriophage (MS2-RFP) coat protein expressed from the viral *nef* locus (Vif+ and Vifxx Gag-CFP/MS2-RFP viruses, see cartoon depictions in Figs. 3A and 3C). When plasmids encoding visible HIV-1 were transfected into cells, Gag-CFP expression allowed for single cell measurement of viral late gene expression, with genomes tracked over time using the MS2-RFP proxy. Expression of Vif+ or Vifxx Gag-CFP/MS2-RFP viruses recapitulated effects seen during infection with CFP reporter viruses (see Fig. 1); including rapid, Vif-dependent down-regulation of YFP-A3G (decay rate = 25.9% per hour +/− 7.0%; n = 31, Fig. 3B, green, with quantification in Figs. 3E and 3F, see also Movie S5) and, for Vif-deficient (Vifxx) conditions, co-clustering of YFP-A3G, MS2-RFP-tagged genomes, and Gag-CFP to assembly sites at the plasma membrane (Fig. 4D, arrows, Movie S6; and see Fig. 4D for an example at higher magnification). Vif-induced YFP-A3G degradation occurred prior to the onset of virus particle assembly (Fig. 3B, compare 2 h and 3 h time points) but >1h after genomes populated the cytoplasm (Fig. 3D, compare 1h and 4h time points). A “Genome only” variant was also generated wherein we mutated the *gag* start codon to abolish Gag synthesis (mutation previously described in ref. (39)) (Fig. 3G). We also generated a version wherein we deleted the Rev response element (RRE) in the second major intron in order to abolish genome nuclear export (see Fig. 5F).

**Figure 3.**
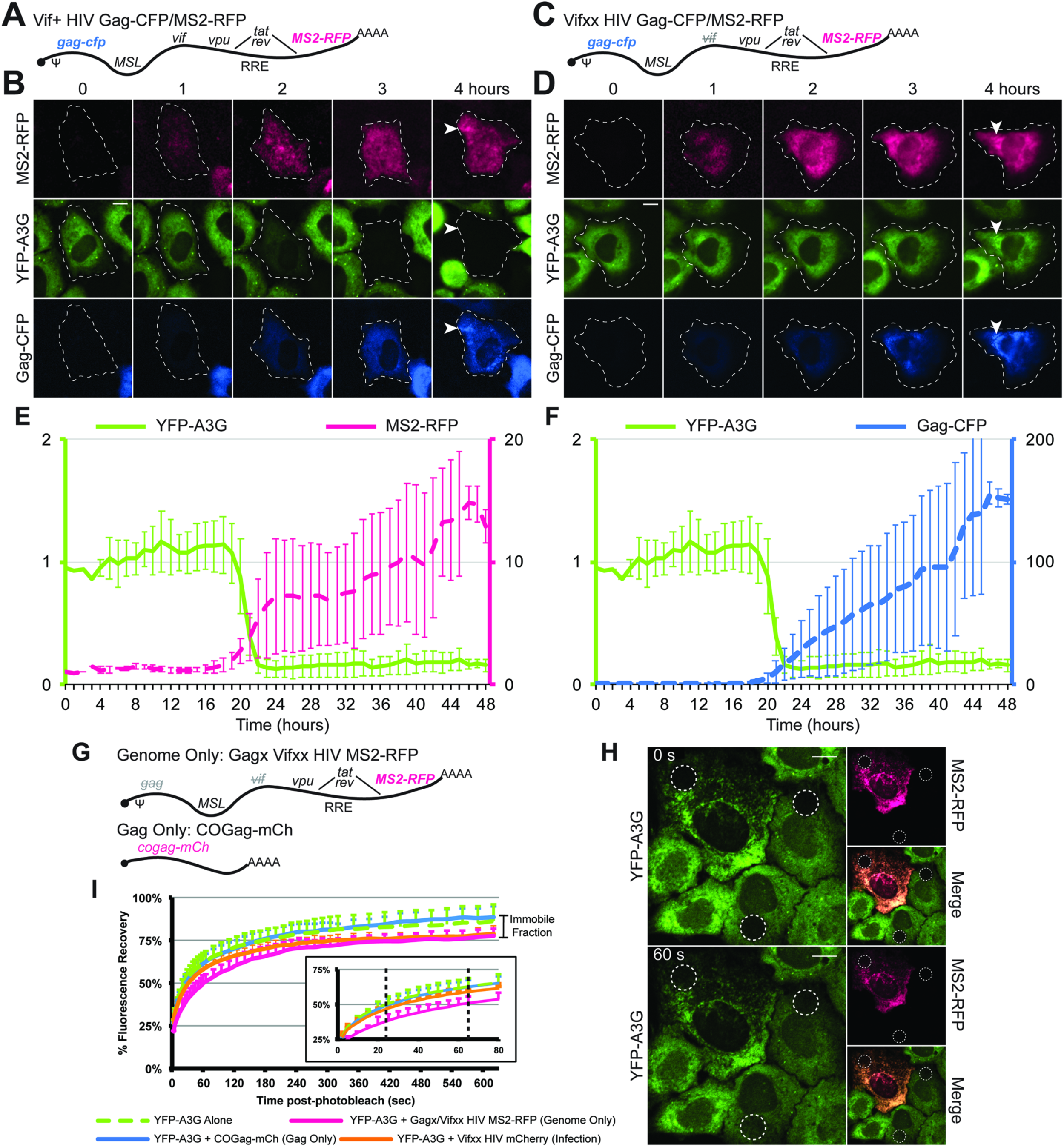
HIV-1 RNA genomes but not Gag regulate A3G subcellular localization. **(A)** and **(C)** Schematics of Vif+ and Vifxx two-color self-tagging viruses encoding Gag-CFP, 24xMSL, and MS2-RFP. **(B)** and **(D)** Time-lapse images of cells expressing Vif+ and Vifxx two-color HIV virus showing nuclear export of MS2-RFP tagged genomes, YFP-A3G degradation (Vif+ in **(B)**) or localization with MS2-RFP (Vifxx in **(D)**), and Gag-CFP expression and accumulation in virus particles at the PM (white arrows). Figure corresponds to Movies S4 and S5. **(E)** and **(F)** Quantification of YFP-A3G degradation and MS2-RFP cytoplasmic accumulation **(E)** and Gag-CFP expression **(F)** in single cells during expression of Vif+ two-color HIV-1. Analysis includes cells that could be tracked for at least 17 h (1 image/hour) extracted from three independent experiments (*n*=31). Black lines highlight the ∼3 h window of YFP-A3G degradation. T=0 represents onset of MS2-RFP expression. **(G)** Schematic of “genome only” and “Gag only” RNA constructs used for fluorescence recovery after photobleaching (FRAP) analysis. **(H)** Representative images from prior to (top) and after (before) photobleaching regions of HeLa.YFP-A3G cells expressing HIV-1 genomes in the absence of Gag (“genome only” condition). White dashed circles represent regions of targeted photobleaching. **(I)** Quantification of YFP-A3G FRAP experiments and zoomed inset of time period over first 80 seconds after photobleaching. Incomplete recovery is best explained by an immobilized fraction of YFP-A3G. All scale bars in fluorescent images = 10µm. Error bars represent standard deviation of the mean.

**Figure 4.**
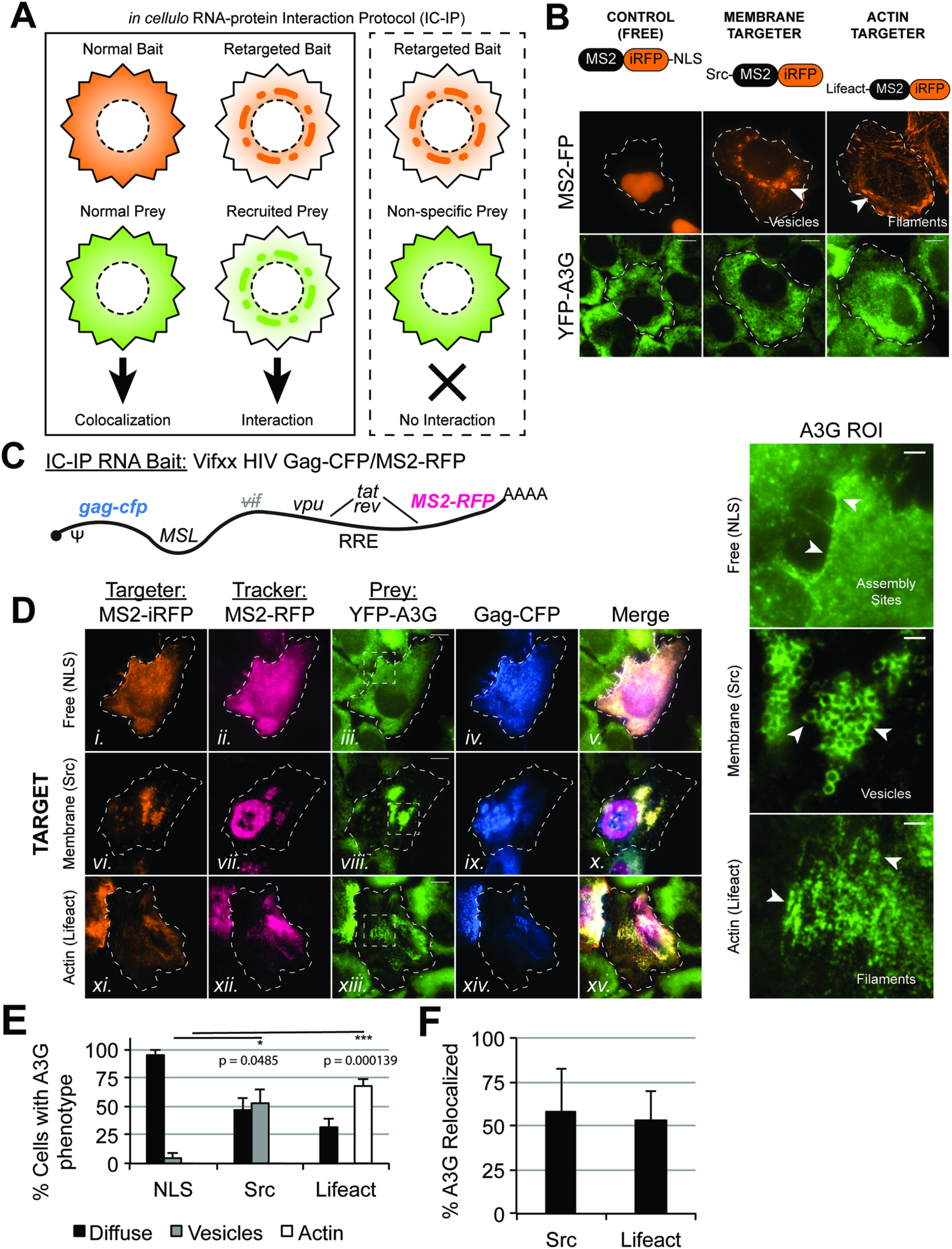
Viral RNA genome-A3G interactions are sufficiently strong to redistribute A3G to unnatural subcellular locales. **(A)** Schematic description of the new <u>I</u>n <u>C</u>ell RNA-protein <u>I</u>nteraction <u>P</u>rotocol (IC-IP). **(B)** Schematic of MS2-iRFP constructs used for “targeting” (top) and representative images of MS2-iRFP and YFP-A3G localization in the absence of mRNAs encoding MSL cassettes. White arrows point to characteristic subcellular targets of Src- (vesicles) or Lifeact- (filamentous actin) MS2-iRFP. **(C)** Schematic of the IC-IP RNA bait used to study YFP-A3G; a Vifxx two-color self-tagging HIV genome. **(D)** Representative images showing “Free” MS2-iRFP-NLS, YFP-A3G, and Gag-CFP accumulating at PM-adjacent assembly sites consistent with the native assembly pathway (top). (Middle) Src-MS2-iRFP accumulating at intracellular/perinuclear vesicles and recruiting MS2-RFP-tagged RNA genomes, YFP-A3G, and Gag-CFP to these vesicles. (Bottom) Lifeact-MS2-iRFP accumulating at F-actin filaments and recruiting MS2-RFP-tagged RNA genomes, YFP-A3G, and Gag-CFP to these F-actin filaments. Regions of interest (ROIs) show YFP-A3G signal in dashed white boxes with arrows highlighting YFP-A3G localization at assembly sites (top) or genome-dependent re-localization to vesicles (middle), or linear F-actin filaments (bottom). **(E)** Single cell quantification of YFP-A3G localization phenotypes (*n = >* 90 cells per condition) in the presence of NLS, Src-, and Lifeact-MS2 targeting constructs. Asterisks demarcate p-values as follows: *≤0.05, **≤0.005, ***≤0.0005. **(F)** Percentages of total YFP-A3G per cell fluorescence (IntDen) re-localized by Src- or Lifeact-MS2 constructs for a subset of cells per condition (*n* > 6). All scale bars in fluorescent images = 10µm, except for YFP-A3G ROIs (1µm). Error bars represent standard deviation of the mean.

**Figure 5.**
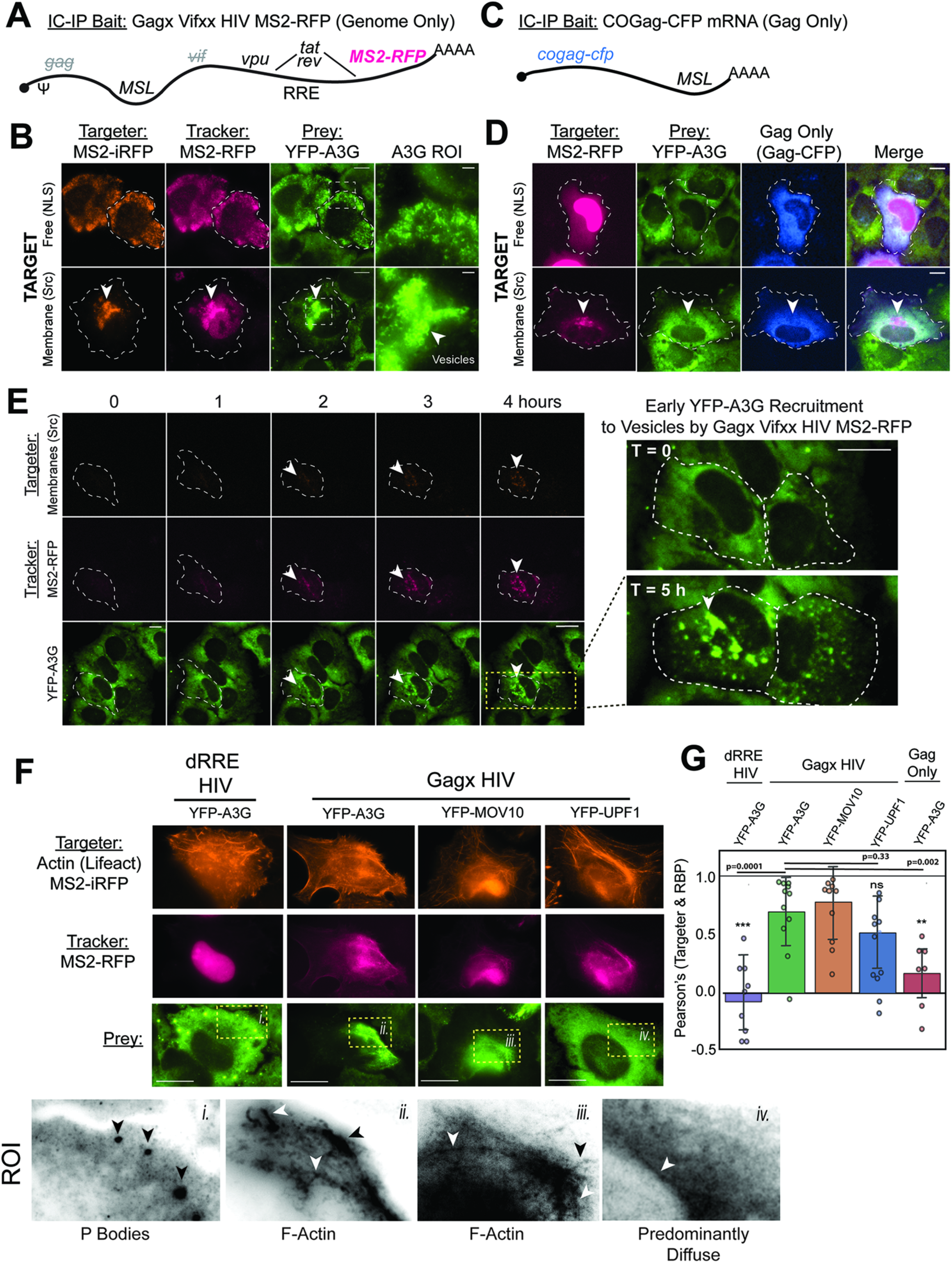
Recruitment of YFP-A3G and MOV10 is selective for HIV-1 RNA genomes, Gag-independent, and occurs long before the onset of virus particle assembly. **(A)** and **(C)** Schematics of “genome only” and “Gag only” IC-IP bait constructs, respectively, used in these experiments. **(B)** Representative images showing co-distribution of MS2-RFP tagged RNA genomes and YFP-A3G under native expression conditions (top) vs. retargeting of YFP-A3G to intracellular vesicles when RNA genomes are tethered to vesicles by Src-MS2-iRFP (bottom). **(D)** Representative images showing specificity, wherein Src-MS2-RFP has no effect on YFP-A3G in the presence of “Gag-only” codon-optimized IC-IP bait mRNAs (white arrow). CFP (blue) channel confirms high-level “Gag-only” mRNA expression based on Gag-CFP synthesis. **(E)** Time-lapse images of cells expressing MS2-RFP-tagged HIV-1 genomes in the absence of Gag (“genome only” construct) and being retargeted by Src-MS2-iRFP over a period of 4 hours. T=0 represents the first detection of the MS2-RFP genome proxy. Arrow shows accumulation of YFP-A3G at Src-MS2-iRFP+ vesicles as early as 2 hours after the onset of MS2-RFP expression. All scale bars in fluorescent images = 10µm. **(F)** Representative IC-IP images of HIV genomes and the indicated RBPs retargeted by Lifeact-MS2-iRFP. ROIs *i-iv* shown below in black and white show re-localization of YFP-A3G and YFP-MOV10 to F-actin filaments by HIV-1 genomes even in the absence of Gag (“genome only). The dRRE control expressed an HIV-1 genome wherein the Rev response element (RRE) is deleted to prevent genome export from the nucleus (see MS2-RFP signal). **(G)** Unbiased quantification of IC-IP interactions based on colocalization analysis of Lifeact-MS2-iRFP targeter and YFP-A3G, YFP-MOV10, or YPF-UPF1 RBP prey for the indicated RNA baits. Non-thresholded Pearson’s coefficients were calculated, comparing targeter & RBP overlap from ROIs for ∼10 cells chosen at random per conditions. Error bars represent standard deviation from the mean. Asterisks demarcate p-values as follows: *≤0.05, **≤0.005, ***≤0.0005.All scale bars in fluorescent images = 10µm.

Both prior to and during virus particle assembly, Vifxx visible genomes and YFP-A3G were co-distributed in a non-localized fashion throughout the cytoplasm, with YFP-A3G but not genome or Gag-CFP accumulating at PBs (Fig. 3D, 2 and 3 h time points post-onset of viral MS2-RFP gene expression). To begin to test if genomes and YFP-A3G were interacting in the cytosolic fluid, we performed fluorescence recovery after photobleaching (FRAP) analysis to measure rates of YFP-A3G recovery with or without genomes and in the presence or absence of Gag (Fig. 3H, and quantification in Fig. 3I); according to the hypothesis that interactions with bulky genomes would slow YFP-A3G movements in the cytoplasm. To establish a baseline, we first infected cells with a Vif-deficient HIV-1 reporter virus (Vifxx HIV mCherry), and confirmed that infection indeed markedly reduced rates of YFP-A3G recovery (t_1/2_ = 34.4 s +/− 5.3 s; *n*=8 cells) relative to the uninfected (YFP-A3G alone) control (t_1/2_ = 27.0 s +/− 13.5 s; *n*=14 cells). Interestingly, we also observed incomplete recovery of YFP-A3G in infected cells (∼75% recovery compared to nearly 90% for untreated control cells Fig. 3I); consistent with the notion that infection actually immobilized a significant fraction of the YFP-A3G pool within the cytoplasm (74). To next dissect the relative contributions of genomes vs. Gag, we compared YFP-A3G recovery after expressing genomes alone (Gagx/Vifxx visible HIV; “Genome Only”) to a “Gag Only” condition wherein we expressed mCherry-tagged Gag from an mRNA modified to lack 5’ and 3’ viral regulatory RNA sequences and codon-optimized to further reduce the potential for contributions of viral *cis*-acting RNA elements located in the *gag* open reading frame (see cartoon depiction in Fig. 3G). The “Genome Only” condition yielded markedly slower YFP-A3G recovery (t_1/2_ = 70.8 s +/− 27.6 s; *n*=6 cells) relative to the no virus control (YFP-A3G Alone) and (COGag-mCh; t_1/2_ = 30 s +/− 8.9 s; *n*=7 cells) (Fig. 3I). Combined, because expression of genomes but not Gag slowed YFP-A3G movements in the cytoplasm based on FRAP analysis, we postulated that genome-YFP-A3G interactions are initiated in the cytoplasm independently of Gag expression and well prior to the onset of virus particle assembly.

### RNA tethering confirms genome-A3G interactions to be selective, Gag-independent, and initiated soon after RNA nuclear export

Because FRAP does not distinguish between direct or indirect genome-A3G interactions, we endeavored to develop a single cell assay based on RNA tethering that would allow us to make more direct measurements of RNA-RBP interactions in the cytoplasm; dubbed the <u>I</u>n-<u>C</u>ell RNA-Protein <u>I</u>nteraction <u>P</u>rotocol (IC-IP) (depicted in Fig. 4A). The IC-IP was based on the principle that, should RNA-protein interactions be sufficiently strong and selective, tethering an RNA of interest to unnatural sites in the cytoplasm (*e.g.*, membranes or the actin cytoskeleton) will shift the subcellular localization of the RBPs that are strongly and persistently associated with that RNA.

For genome-A3G IC-IPs, MSL-bearing genomes served as “bait” for YFP-A3G “prey”, as depicted in Fig. 4A. Genome retargeting was achieved by co-expressing “targeter” MS2 coat proteins modified to tether MSL-tagged genomes to either membranes (Src-MS2; MS2 bearing an N-terminal 10 amino-acid membrane targeting signal) or the actin cytoskeleton (Lifeact-MS2; bearing an N-terminal 17 amino acid F-actin targeting signal) (Fig. 4B). Src-MS2 and Lifeact-MS2 targeters were expressed fused to iRFP670 so that we could co-visualize the targeter with visible 2-color HIV-1 expressing Gag-CFP and genomes labelled with the self-tagging MS2-RFP “tracker” (see cartoon depiction in Fig. 4C).

Co-expression of visible Vifxx Gag-CFP/MS2-RFP genomes as bait with the Src-MS2-iRFP or Lifeact-MS2-iRFP targeters caused moderate re-localization of Gag-CFP proteins from the cytosolic pool to intracellular vesicles (Fig. 4D, compare panels iv. and ix.) or F-actin filaments (Fig. 4D, compare panels iv. to xiv.), respectively; consistent with a subset of Gag proteins co-trafficking with their genome substrate in the cytosol as we have previously shown (39). Effects on YFP-A3G, however, were more striking, with >50% of the net YFP-A3G per cell fluorescent signal relocalized to intracellular vesicles or F-actin filaments (Fig. 4D, compare panels ii., viii., and xiii., and see YFP-A3G regions-of-interest; ROI; with quantification in Figs. 4E and 4F). Consistent with our FRAP analyses (Figs. 3H and 3I), “Genome Only” bait was sufficient to recruit >50% of YFP-A3G to intracellular vesicles as directed by Src-MS2, indicating that genome-A3G cytosolic interactions are widespread and occur independently of Gag (Figs. 5A and 5B). By contrast, tethering a control mRNA expressing codon-optimized version of CFP-tagged Gag (“Gag Only”) had no effect on YFP-A3G cytoplasmic distribution (Figs. 5C and 5D), confirming specificity. Collectively, these results suggested that HIV-1 genome-A3G interactions in the cytoplasm are strong, specific, and occur in the absence of Gag.

A useful feature of the IC-IP is that it can be carried out in live cells and coupled to video microscopy to determine the timing of genome-RBP interactions relative to the onset of RNA nuclear export. Live cell imaging of cells co-expressing Src-MS2-iRFP670 with a “Genome Only” RNA bait demonstrated YFP-A3G almost fully re-localized to vesicles within two hours of the onset of viral gene expression and observations of genomes exiting the nucleus (Fig. 5E, compare 1h and 2h time points, and see Movie S7 for detail). Accordingly, strong genome-A3G interactions are initiated concomitantly with or very soon after genome nuclear export, and these interactions do not require Gag expression.

To further address specificity, we performed similar experiments using Src- or Lifeact-tethered genomes as bait for a variety of alternative co-expressed YFP-tagged RBP preys including YFP-MOV10 and YFP-UPF1 (Figs. 5F and 5G and see Fig. S2 for a compilation of representative images). Of this panel, only YFP-MOV10 exhibited a particularly marked re-localization equivalent to if not more pronounced than YFP-A3G. YFP-UPF1 localization was shifted by genome, but to a lesser extent (Figs. 5F and 5G).

### Tethering A3G to membranes arrests HIV-1 genome trafficking and inhibits virus particle production

We reasoned that if A3G’s interactions with genomes are sufficiently strong, then tethering A3G to membranes would, as a corollary, arrest HIV-1 genome mobility in the cytosol and abolish genome trafficking to sites of virus particle assembly. To test this concept, we carried out the “reciprocal” IC-IP experiment outlined in Fig. 6A, wherein YFP- or CFP-tagged A3G was modified to bind membranes using the same N-terminal Src-derived myristoylation signal used for Src-MS2-directed RNA tethering. These fusion proteins were co-expressed with two-color Vif-deficient visible HIV-1 (Vifxx Gag-CFP/MS2-RFP) (Fig. 6B) or “Genome Only” (Vifxx Gagx/MS2-RFP) genomes (Fig. 6C). As would be anticipated for a strong interaction, Src-FP-A3G expression induced marked clustering of MS2-RFP-tagged genomes to perinuclear vesicles with (Fig. 6B) or without (Fig. 6C) Gag expression, with genomes co-localizing with Src-FP-A3G in both instances. To address the more relevant scenario of natively infected cells, we generated a stable cell line constitutively expressing Src-YFP-A3G (HeLa.Src-YFP-A3G cells) and infected these cells with Vifxx reporter virus (Fig. 6D). As expected, we genomes (and, to a lesser extent, Gag) co-accumulated with Src-YFP-A3G at cytoplasmic vesicles at 24 hpi, detected using combined IF/FISH (Fig. 6D, bottom, white arrows). Co-expression of Src-YFP-A3G or a Src-MS2 control with MSL-tagged 2-color genome (Gag-BFP in this experiment) yielded dose-dependent reduction to cytosolic Gag level and net efficiency of virus particle release (Fig. 6E). Taken together, these experiments demonstrate that A3G-genome interactions in the cytosol are sufficiently strong so that altering A3G trafficking effectively repurposes this antiviral factor as a physical inhibitor of HIV-1 genome trafficking.

**Figure 6.**
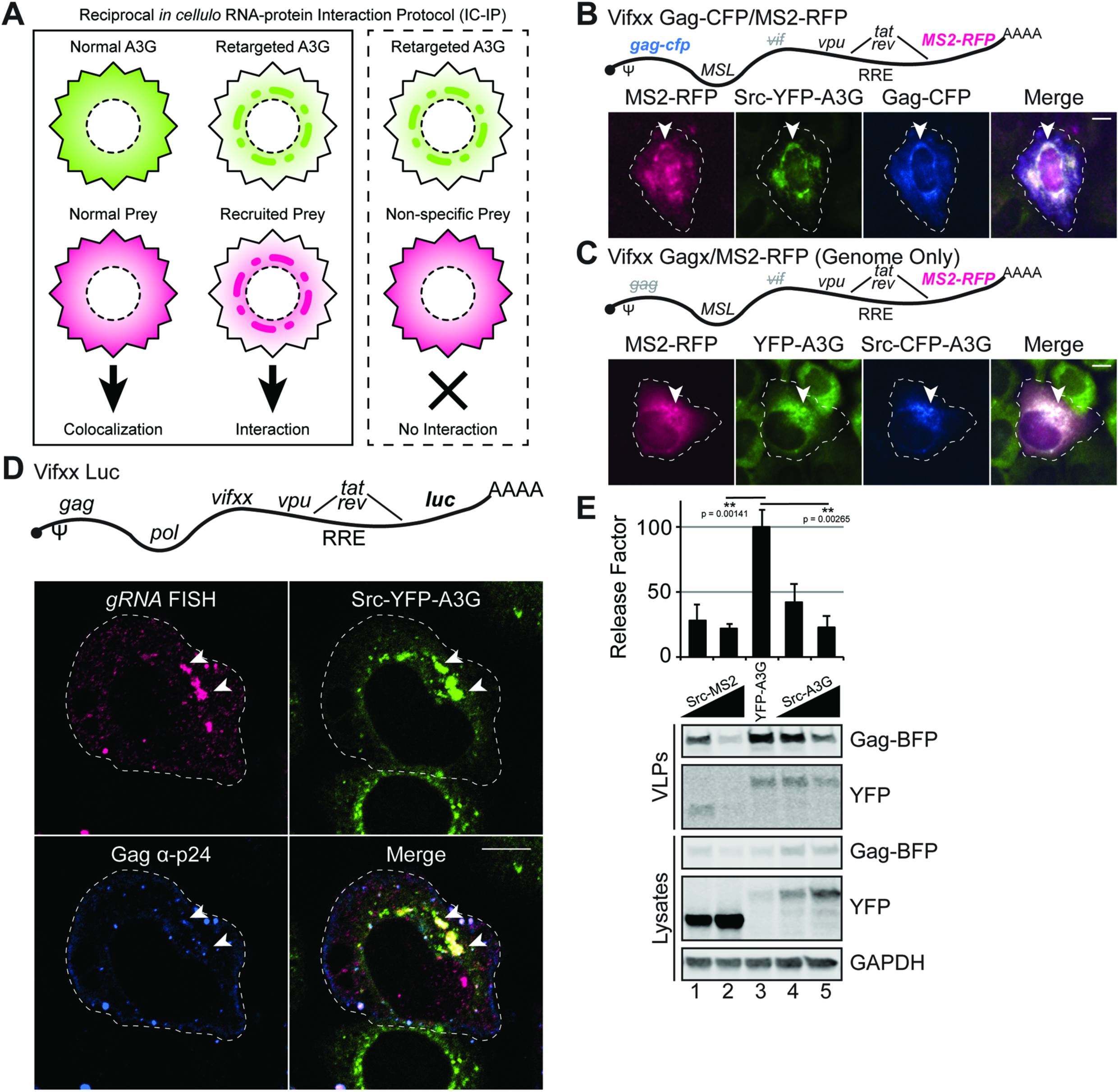
Membrane-targeted APOBEC3G recruits HIV-1 RNA genomes and can inhibit virus particle production. **(A)** Schematic of “Reciprocal IC-IP” experimental set-up to test if Src-YFP-A3G recruits viral genomes to intracellular membranes. **(B)** Schematic of Vifxx 2-color genome and accumulation of MS2-RFP-tagged genomes and Gag-CFP at intracellular vesicles (white arrows) when co-expressed with Src-YFP-A3G. **(C)** As for (B), but using a “Genome only” construct co-expressed with Src-CFP-A3G in YFP-A3G cells. Schematic of Vifxx Luc HIV infectious single-cycle (no MSL) virus used to infect HeLa.Src-YFP-A3G cells and representative images of FISH/IF detecting *gag-pol* mRNA and Gag p-24, respectively. White arrows highlight sites where Src-YFP-A3G recruited *bona fide* HIV RNA genomes to vesicles. **(E)** Western blot showing dose-dependent inhibition of HIV virus particle assembly and production from HEK293T cells transfected to co-express Vifxx two-color HIV-1 with Src-MS2-YFP (lanes 1 and 2), YFP-A3G (lane 3) or Src-YFP-A3G (lanes 4 and 5). Graph shows relative virus particle release efficiency measured as the relative ratio of Gag in virus-like particles (VLPs) to Gag in cellular lysates (“release factor”) quantified from three independent experiments. All scale bars in fluorescent images = 10µm. Error bars represent standard deviation of the mean.

### A3G-genome interactions promote more consistent per virion delivery of A3G to sites of virus particle assembly

Combined, the above experiments suggested that A3G preferentially recognizes one or more HIV-1 genome RNA signatures even in the absence of Gag. However, a remaining conundrum was that A3G is known to be encapsidated by Gag into virus particles even in the absence of packageable genomes (23–25). In an effort to better rationalize prior observations of A3G genome specificity vs. promiscuity, we performed comparative single virion analysis (SVA) of A3G delivery into virus particles using a technique pioneered by the Hu and Pathak groups wherein fluorescently-labeled virus-like particles (VLPs) are harvested from HEK293T cells (that produce a greater quantity of particles relative to HeLa) and subjected to sub-micron quantitative multicolor fluorescence imaging to measure relative levels of per particle genome and/or A3G incorporation (42, 75) (Fig. 7). For this analysis, we compared per particle levels of YFP-A3G for four independent Gag/genome scenarios (depicted in Fig. 7A); (1) “Gag-Only” mRNAs encoding codon-optimized Gag-CFP (COGag-CFP = “Gag-Only”), viral (2) Vif+ or (3) Vifxx 2-color HIV Gag-CFP/MS2-RFP genomes, or (4) mRNAs encoding codon-optimized Gag-CFP wherein the NC RNA-binding domain was replaced by a leucine zipper (ΔNCzip) to abolish Gag-RNA binding (based on (75, 76)). As expected, YFP-A3G packaging was only observed for the COGag and the Vifxx HIV-1 conditions (Fig. 7B).

**Figure 7.**
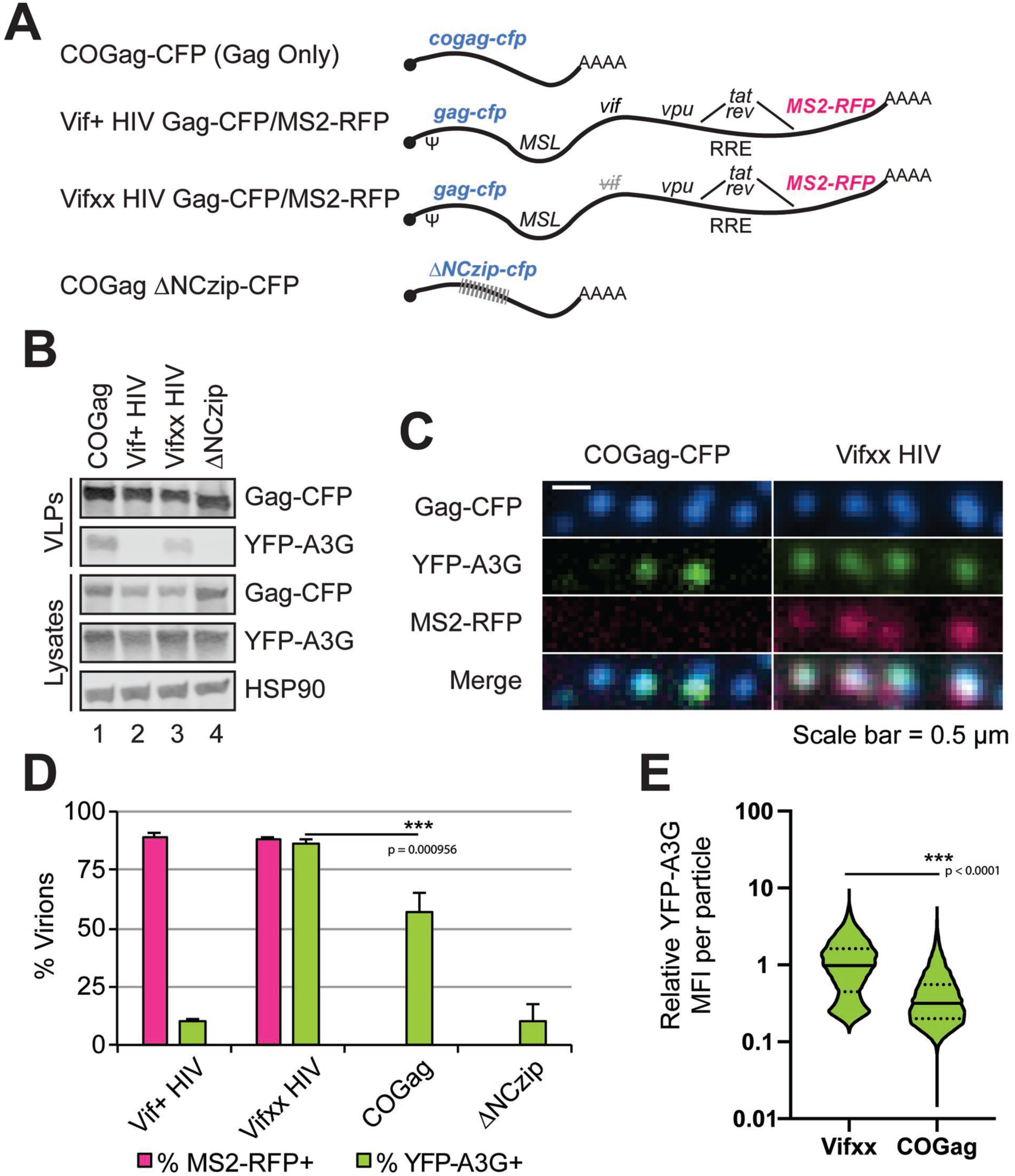
Cytoplasmic A3G-genome interactions promote consistent per virion delivery of A3G into progeny virions. **(A)** Schematics of constructs used for our single virion analysis (SVA) studies. **(B)** Western blot confirming expression and packaging of YFP-A3G only for Vifxx and Gag-only conditions. **(C)** Representative images of single fluorescent virions harvested from our HEK293T.YFP-A3G cell line (see Figure 1C). Scale bar = 500nm. **(D)** Quantification of SVA for each of the four constructs showing percent of virions with MS2-RFP signal (pink) and YFP-A3G (green). Error bars represent standard deviation of the mean (*n* > 13,000 virions per condition). **(E)** Relative YFP-A3G signal per virion or virus-like particle derived from Vifxx two-color or COGag conditions. Relative values are normalized to the mean value for Vifxx with means (solid lines) and 25^th^ and 75^th^ quartiles (dashed lines) indicated.

Using SVA, Vif+ and Vifxx viruses exhibited high efficiency MS2-RFP incorporation (∼89 and ∼88% of Gag-CFP particles scoring positive for MS2-RFP, respectively), consistent with a prior report of HIV-1 genome packaging efficiency (77). Vifxx Gag-CFP/MS2-RFP particles exhibited a similar frequency of YFP-A3G incorporation (86% of total Gag-CFP particles) (Fig. 7C and quantification in 7D). However, for “Gag-Only” (COGag-CFP) particles, the frequency of detectable YFP-A3G incorporation was lower (57% efficiency), intermediate to Vifxx HIV-1 vs. the ΔNCzip “no-RNA” negative control (Fig. 7C and quantification in 7D). COGag particles also exhibited reduced per virion levels of YFP-A3G fluorescence when compared to full-length, Vifxx HIV (Fig. 7C and quantification in 7E). In sum, these data indicated that, although genomes are not essential for YFP-A3G delivery to virus particles, selective genome-A3G interactions promote a more consistent and more enriched per virion delivery of A3G to virus particle assembly sites.

## DISCUSSION

Herein we studied HIV-1 replication from the host cell perspective, using RBP biosensor cell lines, visible HIV-1 viruses, and single-cell RNA-protein interaction assays. On a technical level, our long-term imaging demonstrates that tagged RBP biosensors can be used to detect and measure multivariate aspects of HIV-1 infection. For example, YFP-A3G allows for measurements of Vif-mediated A3G degradation (Fig. 1D and Movie S1), with movies demonstrating Vif’s capacity to degrade A3G is potent, occurs ubiquitously throughout the cell, and is remarkably persistent; lasting hours to days with little to no evidence of fluctuations to Vif activity prior to induction of cell cycle arrest (Figs. 1D, E and Movie S1). We also confirmed that NL4-3 Vif completes its degradation of A3G prior to the onset of virion assembly (Fig. 3B), a result consistent with compelling recent work from Holmes *et al*. who used a similar imaging-based approach to show that Gag’s trafficking to the plasma membrane is delayed until Vif has sufficient time to inactivate A3G (78). Not all Vif alleles degrade A3G (79) or induce cell cycle arrest (80–82). Thus, the A3G biosensor is emerging as a useful tool for categorizing single-cell Vif activities found in circulating HIV strains and subtypes (83).

In the absence of Vif, HeLa.YFP-A3G cells were also able to detect virus particle assembly (Figs. 1F, 1G, S1 and Movies S2, S3), a feat that could also be accomplished using YFP-MOV10 even in the presence of Vif expression (Figs. 1I, S1, and Movie S3). While it was not unexpected that A3G and/or MOV10 would be detected at sites of virus particle assembly, based on prior studies (13, 75, 84), we were surprised to observe such a massive re-distribution of YFP-A3G and YFP-MOV10 into large (assumed to be >2µm diameter at current resolution) clusters of virus particles that remained associated with the cell surface for hours to days, and could be transferred to neighboring cells (see Movies S2 and S3). This clustering phenotype could be attributable, at least in part, to the effects of BST2/Tetherin; a host restriction factor expressed constitutively in HeLa cells that tethers enveloped virus particles to plasma membrane during and after budding (85, 86). However, our viruses encode an intact NL4-3 *vpu* gene that should antagonize BST2/Tetherin. Alternatively, clustering may reflect interactions with charged surface factors such as glycosaminoglycans, that we have previously shown to cause retroviral particle co-clustering and cell retention (“surfacing”) after budding; driven by electrostatic interactions (87). Regardless, to our knowledge this is the first demonstration that HIV-1 assembly and transmission dynamics can be easily recorded using a proxy biosensor (*i.e.*, without having to modify native viruses).

Despite HIV-1’s marked effects on YFP-A3G and YFP-MOV10, we observed no discernible effects of infection on canonical markers of SGs or PBs monitored using our YFP-TIA-1 or CFP-DCP1A biosensors, respectively; with or without Vif expression (Fig. 2). We also observed no major changes to the distribution of NMD factor UPF1 (Fig. 1I, Fig. S1, and Movie S3). These results confirm that HIV-1’s effects on A3G and MOV10 subcellular trafficking are specific; and reinforce that HIV-1 does not modulate visible PBs or induce the formation of SGs, even transiently. The underlying significance of A3G and MOV10’s localization to PBs and SGs remains unknown, but doing so positions these proteins proximal to cytoplasmic mRNA surveillance machines that mediate miRNA-dependent mRNA decay (PBs) or mRNA “triage” during stress responses (SGs) (33–35). In theory, all successful viruses are adapted to subvert, suppress, or avoid such responses; and HIV-1 (88, 89), other retroviruses (90, 91), and endogenous retroelements including LINEs (92) and yeast Ty elements (93) hijack discrete cytoplasmic RNP complexes as a means to compartmentalize key activities including mRNA translation and genome packaging. Regarding suppression, Mouland and colleagues have shown that HIV-1 and other retroviruses actively inhibit SG formation to reduce the potential for cell stress initiated by promiscuous Gag NC binding to host RNAs (69–71). Although we did not observe overt suppression of sodium arsenite-induced SGs by HIV-1 in this study (Fig. 2G-I), the dynamics of these processes over the course of infection warrant further investigation, and it is reasonable to posit that retroviruses must avoid SGs to promote the fluid flow of genomes to capsid assembly sites. Overall, however, our results are most consistent with a model wherein HIV-1’s general *modus operandi* is avoidance of PBs and SGs; with the key interactions underpinning host protein packaging occurring ubiquitously throughout the cytosolic fluid.

Our FRAP and IC-IP results also demonstrate that HIV-1 genomes affect A3G movements in the cytoplasm whether or not Gag capsid proteins are being expressed (Figs. 3-6). Moreover, our data indicate that, in the context of natural infection, A3G (and other RBPs such as MOV10) compete effectively with Gag for genome binding. For example, more than 50% of cytosolic YFP-A3G and MOV-10 were recruited to membranes or F-actin by Src- or Lifeact-MS2-tethered genomes, respectively (Fig. 4F); effects less evident for Gag-CFP, not observed in the absence of genomes (Figs. 5D and 5G), and detected soon if not immediately after genome nuclear export (Fig. 5E). As to *why* A3G-genome binding is selective, single virion analyses (Fig. 7) demonstrated an ∼90% frequency of A3G incorporation into genome-containing virions relative to “Gag-only” particles, wherein the frequency of A3G incorporation was less than 60%. Taken together, these results suggest that A3G combines both promiscuous (25) and targeted (26) RNA binding mechanisms to ensure efficient packaging into HIV-1 virions, with the targeted component enhancing per virion delivery and likely thereby maximizing A3G’s antiviral potential. Regarding the RNA signature detected by A3G, a prior study from York et al. suggested a preference for A3G’s binding to G-rich genome sequences (26), and recent advances in viral epitranscriptomics shows that HIV-1 genomes are enriched relative to host mRNAs in post-transcriptional regulatory marks including N^6^ methyladenosine (m6a) (94–96); and also 5-methylcytosine (m5c) and 2’O-methylation (97). The live cell tools described herein, in particular the IC-IP, should be useful for further dissecting the extent of these interactions as they are occurring in cells.

In sum, our study confirms that the core, essential nature of viral RNA genome trafficking in the cytosol occurs by diffusion in dynamic association with RNP complexes that are smaller than PBs, with some proteins (*e.g.*, A3G and MOV10) more strongly and/or more numerously associated with HIV-1 genomes than others. Because artificially tethering A3G to membranes restricts HIV-1 genome trafficking and virus particle production (Fig. 6), it also remains a possibility that these selective genome binding features could ultimately be exploited in the context of an antiviral strategy.

## MATERIAL AND METHODS

### Cell culture, plasmids, and stable cell lines

Human HeLa and HEK293T cell lines (obtained from ATCC) were cultured in DMEM (Sigma-Aldrich) supplemented with 10% fetal bovine serum (heat-inactivated, filter-sterilized), 1% L-glutamine, and 1% penicillin-streptomycin. HeLa.YFP-A3G, HEK293T.YFP-A3G, HeLa.Src-YFP-A3G, HeLa.YFP-TIA-1, HeLa.CFP-DCP1A, HeLa-YFP-MOV10, and HeLa.YFP-UPF1 cell lines were generated as previously described (39, 43). Briefly, relevant cDNAs were inserted into a MIGR1-derived retroviral vector (pCMS28) (8) upstream of sequence encoding an internal ribosome entry site (IRES) and a second reading frame encoding Puromycin-N-acetyltransferase. Cells were selected by limiting dilution in the presence of 2µg/mL puromycin. YFP, YFP-A3G, and Src-YFP-A3G as well as CFP versions of construct were also generated using the pcDNA3.1 transient expression vector backbone (Invitrogen). RBP expression constructs were cloned similarly into pCMS28, pcDNA3.1, or the pSYFP2-C1 plasmid vector (a gift from Dorus Gadella, Addgene plasmid #22878; http://n2t.net/addgene:22878; RRID:Addgene_22878) using cDNAs generated directly from HeLa cell mRNA (RPL9, or RPS6, CBP80) or derived from the following sources: DCP1A, TIA-1, and PABPC1 (28, 33, 44), MOV10 (a gift of Thomas Tuschl, Addgene plasmid #10977) (45), UPF1 (a gift of Hal Dietz, Addgene plasmid #17708) (46), DDX3x (47), and CRM1/XPO1 (48).

HIV-1 reporter virus plasmids were derived from a modified version of the pNL4-3 X4-tropic HIV-1 molecular clone (49) bearing inactivating mutations in *env*, *vpr*, and expressing a green fluorescent protein (GFP) reporter from the *nef* locus (E-R-/GFP) (50); with the GFP reporter replaced with either cerulean fluorescent protein (CFP) or mCherry (mCh). Vif-minus HIV-1 reporter virus plasmids (Vifxx CFP and Vifxx mCherry) were generated by changing *vif* codons 26 and 27 to stop codons informed by refs. (51, 52). Two-color visible HIV-1 reporter viruses (Gag-CFP/MS2-RFP) were generated by replacing the *gag* reading frame in Vif+ and Vifxx reporter viruses with *gag-CFP* just upstream of sequence encoding 24 copies of the MS2 bacteriophage RNA stem loop (MSL) (53), and inserting sequence encoding the MS2 coat protein fused to a red fluorescence protein (RFP, either mApple or mCherry) and bearing an SV40 nuclear localization signal (MS2-RFP). Gag was alternatively fused to mTagBFP2 for select experiments (*e.g.*, Fig. 6E) in order to avoid cross-detection when immunoblotting for Src-YFP-A3G using anti-GFP/YFP antisera. In all cases, mutated versions of full-length HIV-1 were generated using overlapping PCR as previously described (39). Src-MS2-CFP, Src-MS2-RFP, and Src-MS2-iRFP targeting constructs encoding an amino-terminal membrane targeting signal derived from the Src kinase (MGSSKSKPKD) were generated by overlapping PCR and subcloned into pcDNA3.1. mTagBFP2, mTurquoise2 (CFP), and mApple (RFP) were gifts of Michael Davidson (Addgene plasmids # 54843, 54747, and 55302, respectively) (54, 55). iRFP670 cDNA (56) was amplified from the ColorfulCell expression plasmid, a gift of Pierre Neveu (Addgene plasmid # 62449) (57).

### Retroviral assembly and infectivity assays

Cells at 30-40% confluency were transfected with 2µg DNA in six well plates using polyethylenimine (PEI; #23966, Polysciences Inc.). pcDNA3.1 or pBlueScript were used as empty vector controls. Culture media were replaced at 24 hours post-transfection and cell lysates and supernatants were harvested for immunoblot analysis at 48 hours as previously described (58). Briefly, 1mL of harvested culture supernatant was filtered, overlaid on 20% sucrose (w/v) in PBS, subjected to centrifugation at >21,000g for two hours at 4°C, and viral pellets were resuspended in 35µL dissociation buffer (62.5 mM Tris-HCl, pH 6.8, 10% glycerol, 2% sodium dodecyl sulfate [SDS], 5% β-mercaptoethanol). Cells were harvested in 500µL radioimmunoprecipitation assay (RIPA) lysis buffer (10 mM Tris-HCl, pH 7.5, 150 mM NaCl, 1 mM EDTA, 0.1% SDS, 1% Triton X-100, 1% sodium deoxycholate), homogenized by passage through a 26G needle, subjected to centrifugation at 1,500g for 20 minutes at 4°C, and liquid supernatant fraction was combined 1:1 with 2X dissociation buffer. Proteins were resolved by sodium dodecyl sulfate-polyacrylamide gel electrophoresis (SDS-PAGE) and transferred to 0.2 µm nitrocellulose membranes. Gag was detected using a mouse monoclonal antibody recognizing HIV-1 capsid/p24 (183-H12-5C; 1:1000 dilution) from Dr. Bruce Chesebro and obtained from the NIH AIDS Research and Reference Reagent Program (Bethesda, MD, USA) (59) and anti-mouse secondary antibodies conjugated to an infrared fluorophore (IRDye680LT, 1:10000 dilution, Li-Cor Biosciences) for quantitative immunoblotting. As loading controls, heat shock protein 90A/B (HSP90) was detected using a rabbit polyclonal antibody (H-114, 1:2500 dilution, Santa Cruz Biotechnology, Santa Cruz, CA, USA) or glyceraldehyde 3-phosphate dehydrogenase (GAPDH) was detected using a mouse monoclonal antibody (6C5, 1:2500 dilution, Santa Cruz Biotechnology) and anti-rabbit or anti-mouse secondary secondary antibodies conjugated to an infrared fluorophore (IRDye800CW, 1:7500 dilution, Li-Cor Biosciences). YFP-containing proteins (*e.g.*, YFP-A3G) were detected using a rabbit polyclonal antibody recognizing GFP (FL sc-8334, 1:1000 dilution, Santa Cruz Biotechnology) and anti-rabbit secondary antibodies conjugated to an infrared fluorophore (IRDye800CW). For infectivity assays, supernatants containing single-round infectious HIV-1 virions (pseudo-typed with VSV-G) were filtered and then added to target HeLa cells in the presence of 10µg/mL polybrene. Approximately 36 hours later, target HeLa cells were scanned using a BioTek Cytation5 to detect reporter CFP (455nm/510nm excitation/emission filter) expression following infection.

### Microscopy, immunofluorescence, and fluorescence in situ hybridization (FISH)

Cells were plated in 24-well glass-bottom plates (Eppendorf) or 8-well microslides (IBIDI) and transfected using PEI or infected with HIV-1 reporter viruses. Transfection mixes contained 1µg (24-well) or 333ng (IBIDI) total plasmid DNA, respectively. Infections were performed with VSV-G pseudo-typed HIV-1 reporter viruses produced in HEK293T cells and titered on HeLa cells to determine MOI of 0.5-1, thereby ensuring that ∼33-66% of the cells would be infected per experiment. Fixed-cell experiments were performed on a Nikon Ti-Eclipse inverted wide-field microscope (Nikon Corporation, Melville, NY, USA) using a 100x Plan Apo oil objective (numerical aperture [NA], 1.45). Transfected cells were fixed 24 – 32 hours post-transfection and infected cells were fixed 42 hours post-infection using 4% paraformaldehyde. If treated with 250 µM sodium arsenite (Sigma-Aldrich), drug or an equivalent volume of dimethyl sulfoxide were added to cell culture wells at approximately one hour prior to fixation and returned to incubation at 37°C. Live-cell imaging experiments were also performed on a Nikon Ti-Eclipse inverted wide-field microscope using a 20x Plan Apo objective lens (NA, 0.75) with images acquired every 60 minutes (except where otherwise indicated) over a time course of 16 to 90 hours. Images were acquired using an ORCA-Flash4.0 CMOS camera (Hammatsu Photonics) and the following excitation/emission filter sets (nanometer ranges): BFP (402/455), CFP (430/470), YFP (510/535), mApple (555/605), mCherry (572/632), and iRFP (645/705). All images were processed and analyzed using FIJI/ImageJ2 (60). For IC-IP colocalization analysis, ∼10 targeter single cell iRFP+/RFP+ channel ROIs were chosen at random, used to build ∼1 µM z-stack max intensity projections prior to comparing the overlap of iRFP (targeter) and YFP (RBP) channels using the Coloc2 plugin in FIJI/ImageJ2.

For fixed-cell experiments using FISH, cells were plated as described as above and previously (39). At 42 hours post-infection or after 24 hours plated (for uninfected cells), cells were washed, fixed in 4% formaldehyde, and permeabilized in 70% ethanol for at least 4 hours at 4°C. Custom Stellaris FISH probes were designed to recognize NL4-3 HIV-1 *gag-pol* reading frame nucleotides 386-4456 using the Stellaris RNA FISH Probe Designer 4.1 (Biosearch Technologies, Inc.) available online. To detect *yfp* mRNAs, we used a DesignReady Probe set specific to eGFP. Both probe sets were labeled with CAL Fluor Red 590 Dye (Biosearch Technologies, Inc.). Samples were hybridized with the *gag/gag-pol* probe set according to the manufacturer’s instructions available online. Simultaneous immunofluorescence to detect Gag used a mouse monoclonal antibody specific to p24 (#24-2, a gift of Dr. Michael Malim). Imaging experiments were performed as described above on a Nikon Ti-Eclipse inverted wide-field microscope using a 100x Plan Apo objective lens.

FRAP experiments were performed using a Nikon Ti-Eclipse inverted A1R+ resonant/galvano hybrid confocal line-scanning microscope. Images were captured using a 20x Plan Apo objective lens (NA, 0.75) and GaAsP multi-detector for 488 and 560nm channels. YFP-A3G was imaged using the 488nm laser at a low arbitrary intensity and photobleached using the laser’s maximum intensity. MS2-RFP and COGag-mCherry were imaged using the 560nm laser at a low arbitrary intensity. Cells were maintained in an environmental chamber (Pathology Devices, Inc.) at 37°C, 5% CO_2_, and 50% humidity. Cells were imaged every 30 seconds for 4 frames prior to photobleaching (3 rapid ablations of cytoplasmic ROIs 10µm in diameter) followed by imaging every 5 seconds for a minute, every 15 seconds for four minutes, and every 30 seconds for 5 minutes. This time frame was sufficient for fluorescence recovery to reach a plateau. FRAP analysis was performed using the FIJI plugin FRAP Profiler (61) that adjusts for incidental field photobleaching outside the ROI but within the cell of interest.

Single virion analyses (SVA) were performed largely as described by Chen *et al.* (42). Virus particles were produced as described above for western blotting in HEK293T.YFP-A3G cells to maintain consistent levels of YFP-A3G as compared to co-transfection. Filtered culture supernatants (750µL) were purified by sucrose centrifugation as described above and resuspended in 100µL 1x PBS (Sigma-Aldrich). 24-well plates were pre-coated with 2% FBS diluted in 1x PBS for at least 30 minutes, this solution was removed, and the resuspended concentrated virus particles were added to wells. Images were acquired on a Nikon Ti-Eclipse inverted wide-field microscope using a 100x Plan Apo objective lens (NA, 1.45). These images were processed and analyzed using Analyze Particles plugin in FIJI/ImageJ2 (60).

### Thin-section electron microscopy

HeLa.YFP-A3G were infected with Vif+ CFP or Vifxx CFP HIV-1 viruses and fixed at 48 hours for chemical processing as previously described (62). Samples were sectioned into 100nm slices and with sections collected on copper thin-bar grids. Sections were observed with a Phillips CM120 transmission electron microscope, and images were collected with a MegaView III (Olympus-SIS, Lakewood, CO, USA) side-mounted digital camera. All images were processed and analyzed using FIJI/ImageJ2 (60).

### Statistics

For assembly assays (e.g., Figs. 1C and 6E), results were obtained from three independent biological replicates; defined as cells plated and processed on different days. Graphs plot the mean value with error bars representing the standard deviation of the mean with the exception of Fig. 5G, that shows the second and third quartiles and Fig. 7E, a violin plot showing all data points with the mean (solid line) and quartiles (dashed lines) indicated. All statistical comparisons were carried out using the two-tailed Student’s *t* test and performed using Microsoft Excel or Graphpad Prism.

## ACKNOWLEDGEMENTS

This study was supported by National Institutes of Health [R01AI110221, U54AI150470 to N.M.S., T32CA009135 to E.L.E.III, and T32GM008349 to B.E.B.]; the Wisconsin Partnership Program New Investigator Program [ID 2830 to N.M.S.]; the Greater Milwaukee Foundation’s Shaw Scientist Program [to N.M.S.]; a UW-Madison UW2020 Infrastructure Award [to N.M.S]; a UW—Madison office of the Vice Chancellor of Research and Graduate Education (OVCGRE) Research Competition Award [to N.M.S.]; two National Science Foundation Graduate Research Fellowships [DGE-1256259 to J.T.B. and B.E.B.], an OVCGRE Dissertation Completion Fellowship [to J.T.B.]; an Advance Opportunity Fellowship from the UW-Madison SciMed/GRS program [to E.L.E.III].; and a UW-Madison Hilldale Undergraduate Research Fellowship [to S.L.P.]. Any opinions, findings, and conclusions or recommendations expressed in this material are those of the authors and do not necessarily reflect the views of the National Science Foundation.

We thank Randall Massey at the University of Wisconsin SMPH Electron Microscopy Facility for assistance with EM sample preparation. We are grateful to Michael Malim (King’s College London), Chad Swanson, (King’s College London), and Sarah Gallois-Montbrun (Institut Cochin) for plasmid reagents and advice. The following reagent was obtained through the NIH AIDS Reagent Program, Division of AIDS, NIAID, NIH: HIV-1 p24 hybridoma (183-H12-5C) (from Bruce Chesebro) (59). We thank Kelly Watters, Marchel Hill, and Ann Palmenberg for RVA16.

## SUPPLEMENTAL FIGURE LEGENDS

**Figure S1.**
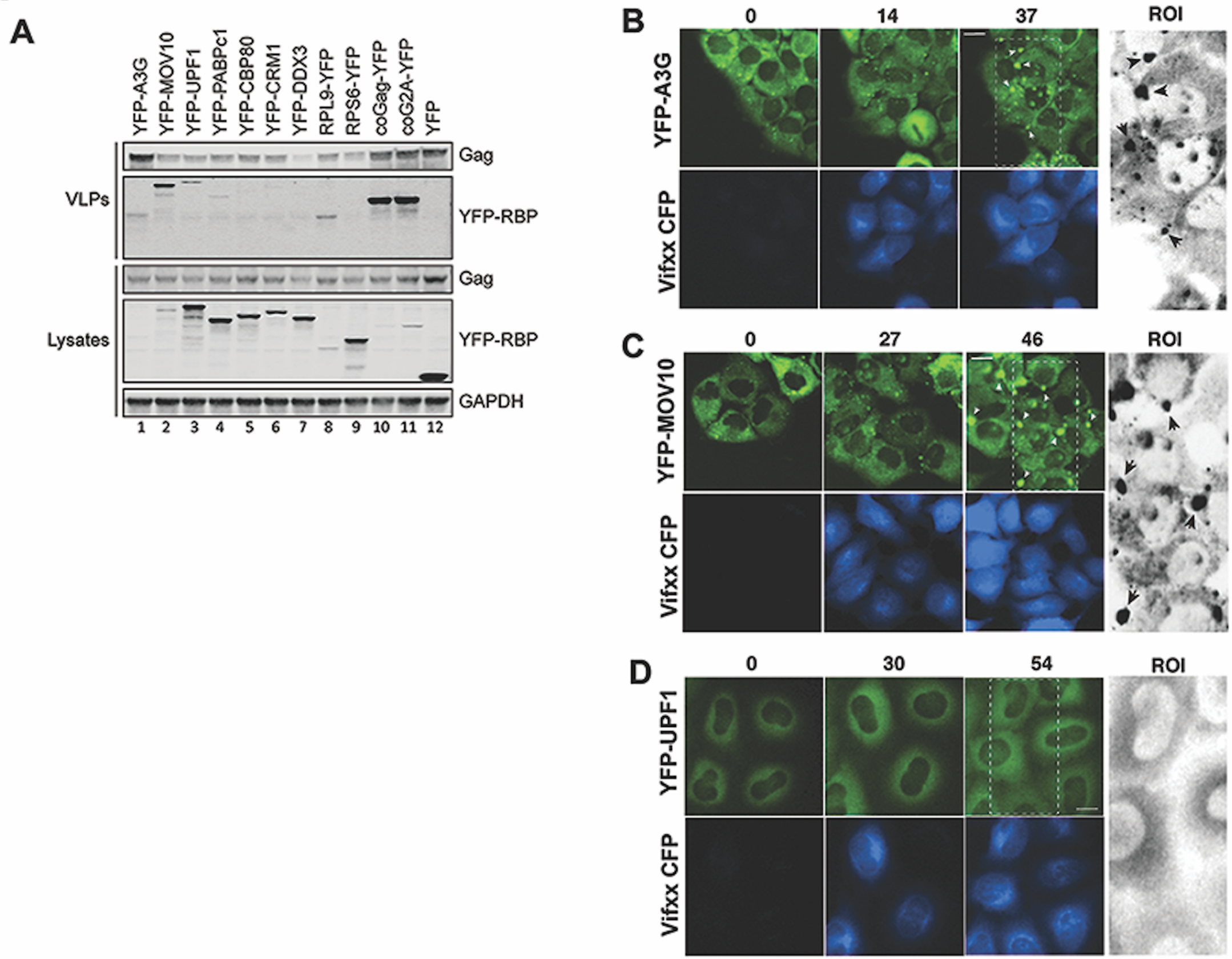
HIV-1 infection alters shifts MOV10 but not UPF1 from the cytoplasm to the plasma membrane. **(A)** Overexpression screen of YFP-tagged RBPs. 293T cells were transfected to generate PR-minus HIV-1 VLPs with co-expression of the indicated YFP-RBP proteins (or YFP as a control, lane 12), with lysates and supernatants harvested at 48 h. GAPDH served as a loading control. (**B-D).** Representative images from time-lapse imaging showing relocalization of YFP-A3G and YFP-MOV10 but not YFP-UPF1 during infection with HIV-1 Vifxx CFP virus. Black and white ROIs highlight >1 µm clusters of YFP-A3G and YFP-MOV10 enriched at the plasma membrane of infected cells at later time points, not observed for YFP-UPF1. **(E)** Select images from time lapse videos showing YFP-MOV10 clusters at the plasma membrane both in the presence and absence of Vif expression. Size bars represent 10 µm.

**Figure S2.**
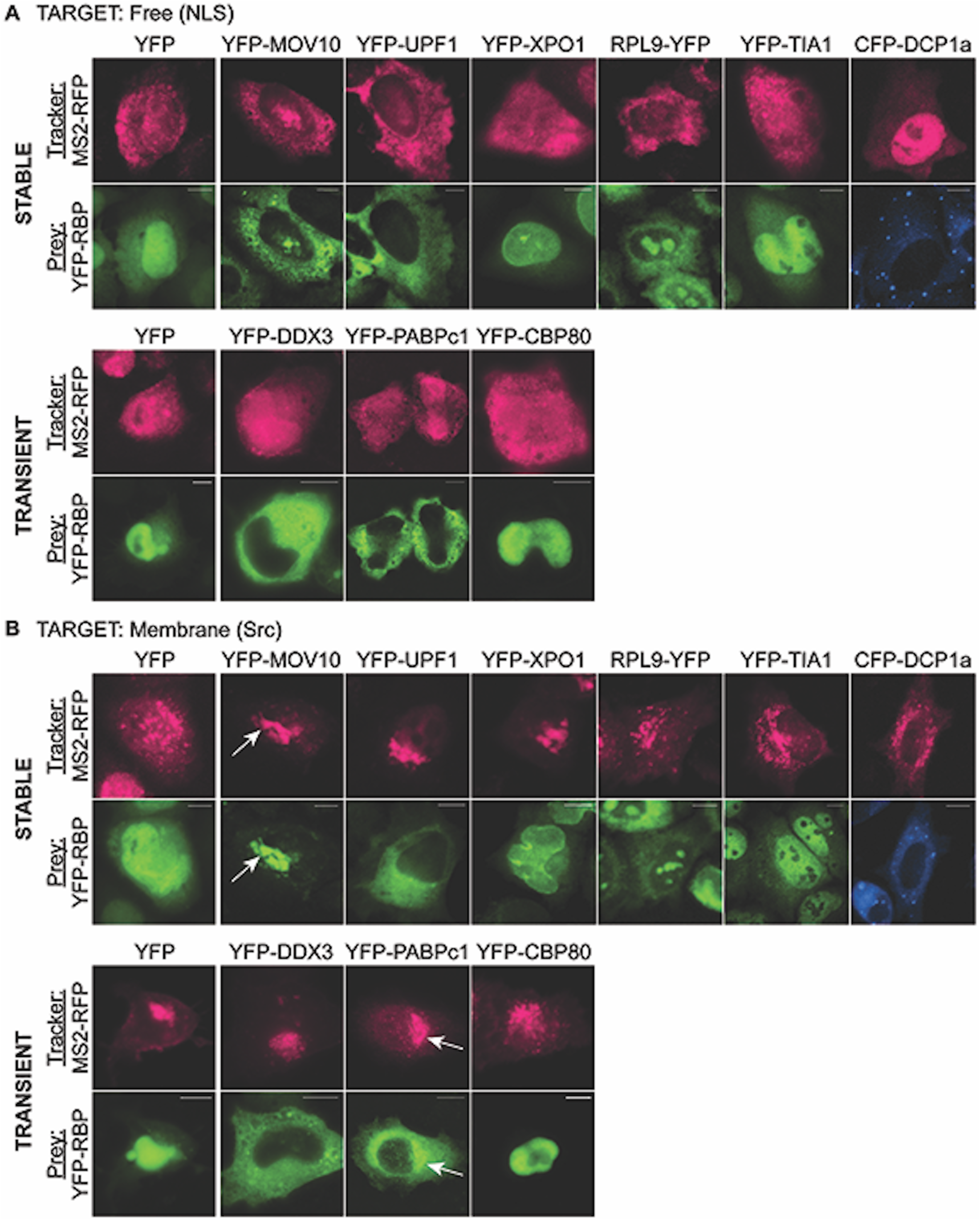
Cellular RBPs are recruited by HIV-1 RNA genomes. (A) Representative images showing co-distribution of MS2-RFP tagged RNA genomes (pink) and YFP-RBPs (green) under native expression conditions. CFP-DCP1A is shown in blue. Top panel shows images from transfections in cells stably expressing YFP-RBPs. Bottom panel shows images from co-transfection of YFP-RBPs with self-tagging HIV genome. (B) Representative images showing co-distribution of MS2-RFP tagged RNA genomes (pink) and YFP-RBPs (green) during retargeting to intracellular vesicles when RNA genomes are tethered to vesicles by Src-MS2-iRFP. Arrows indicate intense recruitment of MOV10 and (to a lesser degree) PABPc1. Other RBPs show minimal (XPO1, UPF1, DDX3) or no (RPL9, TIA-1, DCP1A, CBP80) recruitment to Src-MS2-iRFP retargeted HIV genomes.

## SUPPLEMENTAL MOVIE LEGENDS

**Movie S1. Time-lapse imaging of HeLa-.YFP-A3G cells infected with Vif+ HIV-1/CFP virus.** Multi-channel images of HeLa.YFP-A3G cells were acquired every 60 minutes for 49h after infection. In the highlighted cell, viral gene expression (cyan) is detected ∼16hpi with YFP-A3G (green) degradation at 21hpi. Cell rounds at 28 hpi consistent with Vif-induced G2/M cell cycle arrest.

**Movie S2. Time-lapse imaging of HeLa.YFP-A3G cells infected with Vifxx HIV-1/CFP virus.** Multi-channel images were acquired every 60 minutes for 49h after infection. In the highlighted cell, viral gene expression (cyan) is detected ∼19hpi with detection of YFP-A3G (green) at virus particles (arrows) starting at ∼25hpi. An example of a transfer event wherein a large cluster of YFP-A3G+ virus particles is released from an infected cell is highlighted (arrows) at 32-36hpi.

**Movie S3. Time-lapse imaging of HeLa.YFP-MOV10 and HeLa-YFP-UPF1 cells infected with Vifxx HIV-1/CFP virus.** Multi-channel images were acquired every 60 minutes for 96hpi with Vif+ or Vifxx HIV-1/CFP viruses. A 72h time window is shown, initiating ∼3 hours prior to the onset of viral gene expression (cyan). Vif+ HIV-1 degrades YFP-A3G (top left panel, green) while Vifxx HIV-1 triggers relocalization of YFP-A3G (bottom left panel, green) to plasma membrane clusters (arrows). YFP-MOV10 (central panels, green) is re-localized to plasma membrane clusters (arrows) with or without Vif expression. YFP-UPF1 was largely unaffected by HIV-1 in the presence or absence of Vif, but with some rare instances of cytoplasmic granule formation.

**Movie S4. Time-lapse imaging of HeLa.YFP-TIA-1 cells infected with Rhinovirus A16.** Multi-channel images were acquired every 30 minutes after addition of RVA16 (MOI∼10). The first instances of SG formation (YFP-TIA-1 in green) are evident at ∼11hpi.

**Movie S5. Time-lapse imaging of HeLa.YFP-A3G cells expressing the two-color Vif+ Gag-CFP/MS2-RFP HIV-1 reporter virus.** Multi-channel images were acquired every 60 minutes for 24h beginning ∼4h post-transfection. Left panel shows genome tracked using MS2-RFP (magenta), with nuclear export detected at T=4h just prior to the onset of Gag-CFP (cyan) expression (right panel) and YFP-A3G (green) degradation (center panel).

**Movie S6. Time-lapse imaging of HeLa.YFP-A3G cells expressing the two-color Vifxx Gag-CFP/MS2-RFP HIV-1 reporter virus.** Multi-channel images were acquired every 60 minutes for 24 hours beginning ∼4 hours post-transfection. Left panel shows genome tracked using MS2-RFP (magenta), with nuclear export detected at T=6h just prior to the onset of Gag-CFP (cyan) expression (right panel) and co-accumulation of YFP-A3G (green, center panel) with genome (MS2-RFP) and Gag-CFP at the plasma membrane (at putative sites of virus particle assembly).

**Movie S7. Time-lapse imaging of HeLa.YFP-A3G cells co-expressing “genome only” MS2-RFP HIV-1 reporter virus with the Src-MS2-iRFP targeter (orange).** Multi-channel images were acquired every 60 minutes for 15h starting ∼4h post-transfection. MS2-RFP tracked genomes (magenta, central panel) are first observed at T=0, with both genomes and YFP-A3G (green, right panel) immediately recruited to perinuclear vesicles, co-localizing with the Src-MS2-iRFP targeter (yellow, left panel). Video demonstrates that genomes have marked effects on YFP-A3G trafficking in the cytosol even at the earliest time points post-genome nuclear export.

## Notes

### Competing Interest Statement

The authors have declared no competing interest.

